# Agent_K: An Agent-Orchestrated Pipeline for Mapping Genes to Biosynthetic Pathway towards Metabolite Discovery

**DOI:** 10.64898/2026.10.19.760272

**Authors:** Raj S. Roy, Jeong Jae Han, Jayaram Lakshmaiah Narayana, Aifen Zhou, Paul de Figueiredo, Arum Han, Xiaoning Qian

## Abstract

The identification of biosynthetic gene clusters (BGCs) and their associated metabolic pathways is central to genome-guided natural product discovery. However, existing computational tools depend on fragmented workflows for data processing, annotation, and analysis, which require substantial manual intervention and lack seamless integration across heterogeneous platforms and tools. Here, we present, Agent_K, an agent-orchestrated framework that automates the end-to-end process from bacterial whole genome sequence analysis to biosynthetic pathway mapping and metabolite discovery as candidate gene products for functional insights. The system employs a large-language-model (LLM) agent to coordinate multiple bioinformatics tools, enabling dynamic execution, data integration, result interpretation, and flexible and intuitive visualization. By integrating genomic sequences to functional annotations and pathway-level reconstructions, the framework facilitates the identification of candidate gene products, their biosynthetic pathways associated with individual BGCs, and the putative metabolites produced by those pathways. The developed novel agentic framework provides a user-friendly scalable and extensible solution for automated high-throughput analysis and mapping of bacterial secondary metabolism pathways for interactive exploration of candidate potential.

## 1 Introduction

Natural product discovery relies on uncovering biosynthetic gene clusters (BGCs) that encode enzymes responsible for producing biologically active secondary metabolites. These metabolites, including antibiotics, toxins, and signaling molecules, often arise from complex, multi-step enzymatic pathways that are not immediately apparent from genomic data alone. Advances in genome sequencing and high-throughput omics-profiling have significantly facilitated large-scale identification of BGCs [1, 2]. However, translating these predictions into functional insights remains challenging due to the fragmented nature of existing computational tools and the need for manual integration across multiple analysis stages.

For reliable functional annotation that maps genome sequences to corresponding biosynthetic metabolites, traditional pathway reconstruction approaches typically require sequential execution of specialized tools for BGC detection, functional annotation, and pathway mapping. Such workflows are often rigid, difficult to reproduce, and require substantial user intervention. Recent advances in artificial intelligence (AI), especially in large language models and other foundation models [3, 4], have led to the emergence of agent-based systems capable of orchestrating complex, multi-step tasks through dynamic planning and tool integration. These systems have been successfully applied across diverse domains, including software engineering, data analysis, and scientific workflows, where they enable automated execution of tasks that traditionally require manual coordination [5, 6]. By interpreting high-level user queries and decomposing them into structured actions, agentic frameworks can flexibly integrate heterogeneous tools, manage intermediate states, and iteratively refine outputs. This paradigm is particularly well-suited for bioinformatics workflows, where analyses often involve multiple specialized tools with inter-dependent inputs and outputs. For example, the Virtual Lab [6] framework demonstrates the potential of collaborative agentic systems for scientific discovery, where a multi-agent pipeline enabled the design of de novo nanobodies, several of which exhibited strong binding affinity to variants of SARS-CoV-2.

To systematically identify BGCs and characterize their functionalities by searching for corresponding biosynthetic pathways, we develop, Agent_K, a novel agent-orchestrated pipeline that connects genomic data to biochemical function through coordinated tool execution and data integration. In this framework, the agent serves as a central controller, managing tool execution, tracking intermediate data, and integrating outputs into a coherent pathway-level interpretation. This design transforms a conventional linear pipeline into a flexible and interactive system capable of adapting to user inputs and supporting end-to-end biosynthetic analysis. Altogether, this integrative approach enables rapid prioritization of biosynthetic gene clusters with high potential for novel or functionally relevant natural products

## 2 Materials & Methods

### 2.1 System Overview

The proposed framework consists of an agent-based orchestration layer built on a modular collection of bioinformatics tools. The agent coordinates task planning, tool execution, result integration, and user interaction. By integrating genomic data with information from functional annotation databases, enzyme prediction methods, and metabolic pathway repositories, the framework links the biosynthetic genes to its biochemical function. This enables the identification of enzymes and reactions encoded within a BGC, reconstruction of putative biosynthetic pathways, and prediction of candidate metabolites along with their potential synthesis routes.

### 2.2 Agent-Orchestrated Workflow

Unlike traditional static pipelines, the proposed agentic framework dynamically coordinates the execution of multiple analysis modules based on the user’s demand, leveraging a large language model (LLM) agent, which selects the tools and organize into a workflow to execute. This agent’s orchestration layer integrates intermediate outputs to ensure consistency and completeness of the final results. The overall pipeline is shown in **Figure 1**, consisting of the following stages:

**Figure 1:**
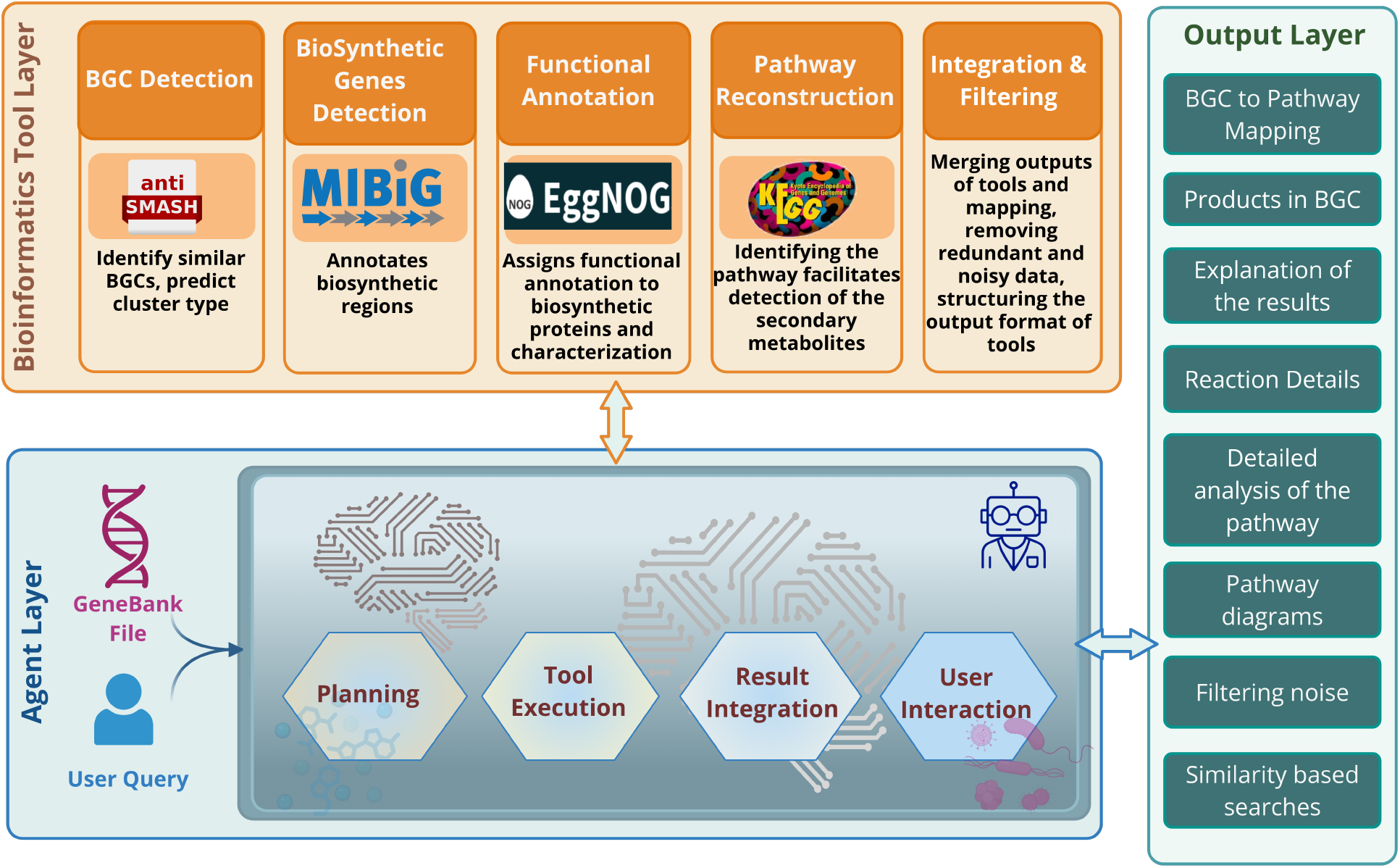
Overview of the agent-orchestrated pipeline, Agent_K, showing the interaction between the central agent, integrated bioinformatics tools, and resulting outputs. The agent coordinates modules for BGC prediction, functional annotation, and pathway reconstruction, and integrates intermediate results to produce pathway-level interpretations and candidate metabolite predictions.

#### 2.2.1 BGC Prediction with AntiSMASH

The agent initiates the analysis by invoking antibiotics & Secondary Metabolite Analysis Shell (AntiSMASH) [7] to identify biosynthetic gene clusters from genomic input sequences. AntiSMASH detects characteristic biosynthetic domains and annotates cluster boundaries and product classes. This step produces annotated BGCs with gene composition, product type, and similarity to characterized clusters.

#### 2.2.2 Biosynthetic Genes Prediction with MIBiG

To contextualize predicted clusters, the user may ask the agent to extract the additional details of the BGC. The agent then extracts additional details of the corresponding cluster. To do so, the agent leverages the Minimum Information about a Biosynthetic Gene cluster (MIBiG) [8] file to extract the “biosynthetic related” protein sequences for downstream functional analysis.

#### 2.2.3 Functional Annotation with EggNOG-mapper

The agent submits these extracted protein sequences to evolutionary genealogy of genes: Non-supervised Orthologous Groups (EggNOG-mapper) [9] for functional annotation. This step assigns orthologous groups and predicts enzyme functions, reactions, and pathway associations. This annotated protein with functional roles and Kyoto Encyclopedia of Genes and Genome (KEGG) [10] ortholog identifiers lays the foundation of pathway identification. This step outputs details of the reactions, products, and pathways, along with structured data that are further filtered and utilized in subsequent steps.

#### 2.2.4 Pathway Reconstruction using KEGG

Using the assigned ortholog identifiers, the agent queries employ a frequency based algorithm to display the most possible pathways. It also displays the alternative pathways as it may be useful for other analysis. Reaction-level and pathway-level mappings, including metabolite information and pathway structures along with other details are produced in this step. Furthermore, the agent makes sure that the global overview of KEGG [10] maps are filtered.

#### 2.2.5 Integration and Filtering

The agent aggregates outputs from all modules into unified data structures. Consolidate and refine the results to filter redundant or low-confidence results, and non-specific pathway mappings are excluded to retain biologically relevant pathways.

### 2.3 Agent Planning and Interaction

The LLM agent performs lightweight reasoning primarily through task decomposition and context-aware tool selection. Upon receiving a user query, the agent interprets the intent and maps it to a sequence of executable actions, such as invoking specific bioinformatics tools or analysis scripts. Rather than relying on a fixed pipeline, the system dynamically selects modules based on the requested task and available intermediate data.

The agent is accessed through a Web-based interface developed using Streamlit and the Qwen2.5-7B [11] base model, a transformer-based large language model with approximately 7 billion parameters. The model is designed to support natural language understanding, instruction following, reasoning, and text generation. Users interact with the system through natural language prompts, which are translated into executable analysis plans. The interface supports iterative exploration of BGCs, pathways, enzymes, metabolites, and associated biochemical reactions while providing real-time access to intermediate and final outputs. By exposing both the execution process and generated results, the system improves transparency, interpretability, reproducibility and speed of analysis compared with conventional command-line workflows.

## 3 Results

GenBank (*.gbk) files derived of the whole-genome sequences of a bacterial strain *Stenotrophomonas sp*. served as the input for genome mining. The strain was selected based on its superior antimicrobial activity against the target drug-resistant pathogen *Pseudomonas aeruginosa* in a high-throughput microfluidics screening platform [12]. Using Agent_K, putative biosynthetic gene clusters (BGCs) were identified and mapped to their associated secondary metabolite biosynthetic pathways to uncover promising candidate natural products. The complete conversation of the user with the agent is illustrated in **Figure 2**. For example, the BGC cluster BGC000343, multiple genes involved in biosynthesis of siderophore, including isochorismate, 2,3-Dihydro-2,3-dihydroxybenzoate, and enterochelin, were mapped to the biosynthetic pathway. Experimental evidence reported independently by our collaborators in a separate study [12] confirmed the production of enterochelin, demonstrating that the pipeline identified the correct pathway associated with it (map01053). Beyond this validated result, the pipeline identified additional candidate pathways, providing testable hypotheses for future experimental studies. To summarize, a total of 46 biosynthetic gene clusters (BGCs) were successfully mapped to reference BGCs and functionally characterized through automated pathway reconstruction. In addition, it identified 18 unique KEGG pathways, 91 unique biochemical reactions, and 94 unique biochemical products. The complete functional annotations, including pathway, reaction, and product information for each mapped BGC, are available in the accompanying GitHub repository.

**Figure 2:**
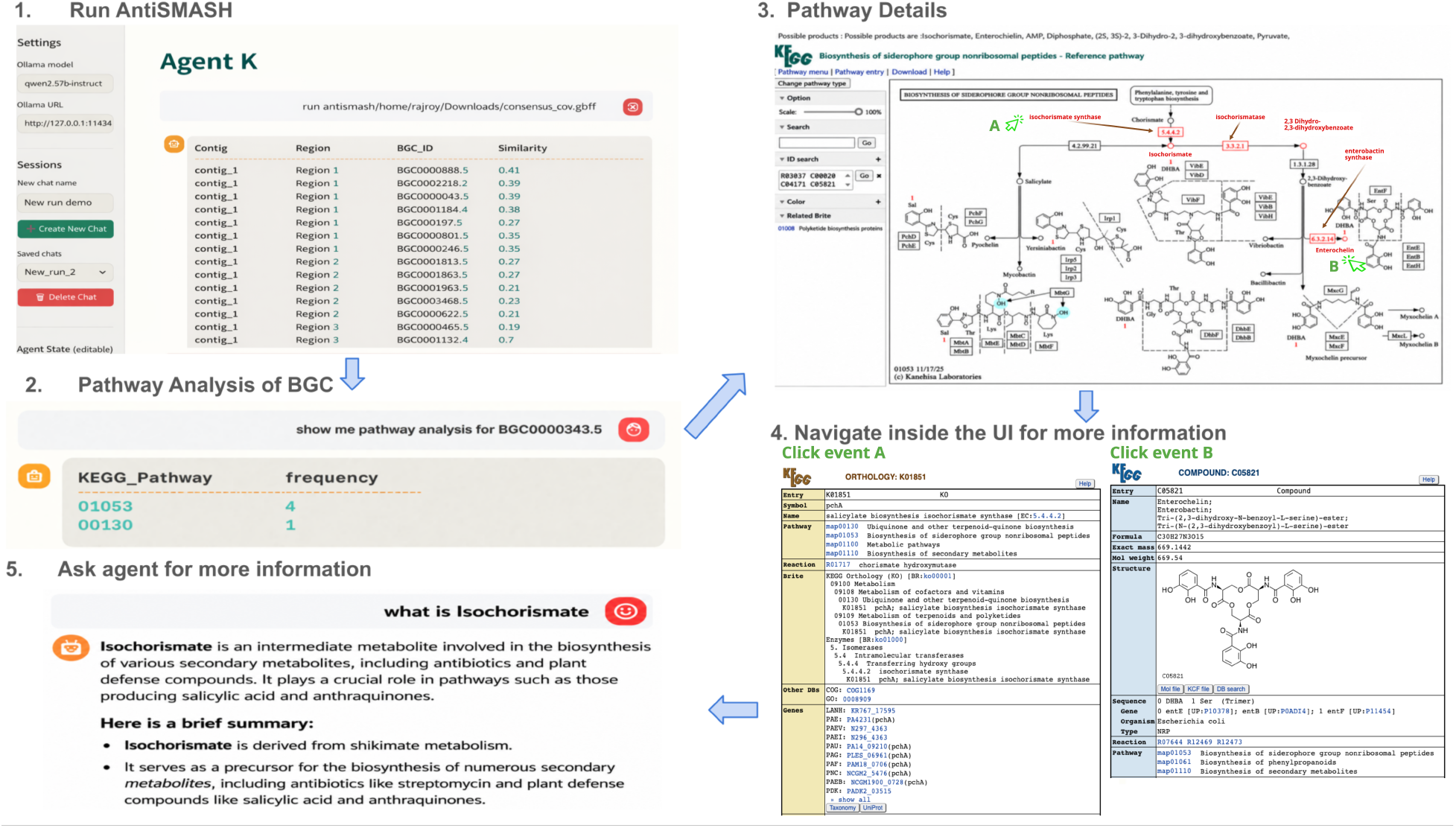
The user interaction with the agent and the corresponding outputs and visualizations. (1) The user provides the respective GenBank file, and the agent identifies candidate biosynthetic gene clusters (BGCs) using antiSMASH. (2) The user selects the BGC0000343.5 for further analysis, and the agent retrieves its associated KEGG pathways. (3) The agent generates annotated pathway visualizations highlighting enzymes, reactions, and products. (4) The user interactively explores pathway components, including compounds and enzymes by clicking the icons in the map. Selected components are highlighted in green in the preceding pathway visualization. (5) The agent integrates pathway results with domain knowledge to provide contextual information about predicted products.

Furthermore, interaction with the agent triggers the execution of multiple tools and scripts, producing intermediate and final outputs at each step of the workflow. These outputs are systematically organized and stored in a structured directory hierarchy, as illustrated in **Figure 3**.

**Figure 3:**
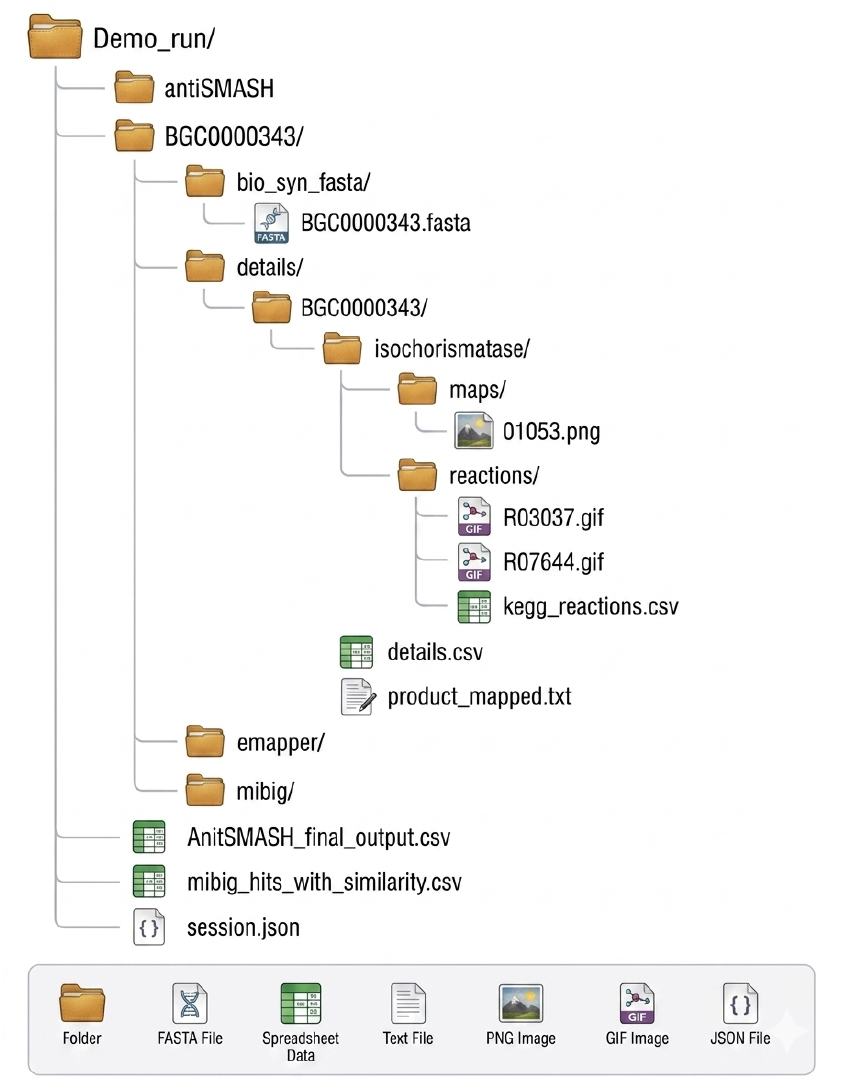
The organized file structure of the intermediate and final output generated during the interactive analysis.

Specifically, outputs from individual tools are grouped into dedicated directories. For example, results from antiSMASH, eggNOG, and MIBiG searches are stored in the directories “antiSMASH”, “emapper”, and “mibig”, respectively. In addition to tool-specific outputs, the agent performs downstream analyses such as removing and filtering products and reactions, and stores the resulting intermediate data in a separate details directory.

The directory of “details” contains processed outputs that facilitate comprehensive interpretation and further analysis. These include pathway visualizations, reaction maps, and tabular summaries of reactants and products. Key files such as kegg reactions.csv and details.csv provide structured representations of the analysis, enabling efficient downstream processing and reproducibility. This organized output design allows users to trace intermediate steps and revisit the analysis workflow when required. Notably, the complete workflow is executed within minutes, details provided in Supplementary Section 1, in contrast to traditional fragmented workflows, which can require weeks of manual data collection, integration, and analysis.

## 4 Discussion

The proposed framework provides a comprehensive bridge from genomic data to pathway-level interpretation, enabling the identification of novel or divergent metabolites, comparative analysis across microbial genomes, and rapid visualization of biosynthetic potential in newly sequenced organisms. By integrating multiple analysis steps into a unified system, it establishes the first reliable end-to-end platform for high-throughput annotation, pathway reconstruction, and exploration of secondary metabolism. In addition, the interactive and agent-driven design facilitates efficient hypothesis generation, reducing the need for manual coordination across disparate tools. Despite these advantages, several limitations should be noted. For instance, while the interface enables interactive exploration, additional user feedback is necessary to optimize usability for a wider scientific audience.

From a methodological perspective, the agent employs a lightweight reasoning strategy primarily focused on task decomposition and tool selection. Although effective for orchestrating structured workflows, this approach could be extended through the incorporation of more advanced agent architectures. For example, multi-agent systems could be introduced to enable specialized roles, such as quality assessment, result validation, or deeper biological interpretation. Finally, deployment as a web-based platform would further improve accessibility and facilitate adoption by the broader research community.

## 5 Conclusion

In this work, we presented a novel agent-orchestrated framework for automated analysis of biosynthetic gene clusters and their associated metabolic pathways. By integrating multiple bioinformatics tools within a unified and interactive system, the proposed approach enables dynamic, end-to-end mapping from genomic sequences to pathway-level interpretations and candidate metabolites. The agent-driven architecture simplifies traditionally fragmented workflows, enhances reproducibility, and supports iterative exploration through natural language interaction.

Overall, this framework provides a scalable and extensible solution for high-throughput analysis of secondary metabolism and facilitates efficient exploration of biosynthetic potential. These results highlight the potential of agent-based systems to streamline complex bioinformatics workflows and accelerate computational discovery in natural product research.

## Code Availability

The genomic input data used in this study were provided by collaborators and are not publicly available due to proprietary data restrictions.

The source code for the Agent_k is publicly available at: https://github.com/RajShekhorRoy/Agent_K, https://doi.org/10.5281/zenodo.23421236

## 6 Competing interests

No competing interest is declared.

## 7 Author contributions statement

R.S.R., A.H., P.D.F., and X.Q. conceived the experiment(s), R.S.R. conducted the experiment(s). R.S.R., J.J.H., J.L.N., A.Z. and X.Q. analyzed the results. R.S.R., and X.Q. wrote the manuscript. A.H., and P.D.F. provided guidance and feedback regarding the pipeline. All authors contributed to the discussions, reviewed and edited the manuscript.

## 8 Acknowledgments

Funding was provided by the National Institutes of Health (NIH), National Institute of Allergy and Infectious Diseases (NIAID), grant #1R01AI168685-01A1, as well as the Texas A&M University Advancing Discovery to Marketplace (ADM) Program grant #292394.

## Notes

### Competing Interest Statement

The authors have declared no competing interest.

